# Exploring the role of life history traits and introduction effort in understanding invasion success in mammals: A case study of Barbary ground squirrels

**DOI:** 10.1101/2020.10.27.357319

**Authors:** Annemarie van der Marel, Jane M. Waterman, Marta López-Darias

## Abstract

Invasive species –species that have successfully overcome the barriers of transport, introduction, establishment, and spread– are a risk to biodiversity and ecosystem function. Introduction effort is one of the main factors underlying invasion success, but life history traits are also important as they influence population growth. In this contribution, we first investigated life history traits of the Barbary ground squirrel, *Atlantoxerus getulus*, a species with a very low introduction effort. We then studied if their invasion success was due to a very fast life history profile by comparing their life history traits to those of other successful invasive mammals. Next, we examined whether the number of founders and/or a fast life history influences the invasion success of squirrels. Barbary ground squirrels were on the fast end of the “fast-slow continuum”, but their life history was not the only contributing factor to their invasion success, as the life history profile is comparable to other invasive species that do not have such a low introduction effort. We also found that neither life history traits nor the number of founders explained the invasion success of introduced squirrels in general. These results contradict the concept that introduction effort is the main factor explaining invasion success, especially in squirrels. Instead, we argue that invasion success can be influenced by multiple aspects of the new habitat or the biology of the introduced species.

## Introduction

Invasive species are a major threat to biodiversity and ecosystem functions, and are an economic concern (Simberloff et al. 2013; Doherty et al. 2016; Gallardo et al. 2016; Bellard et al. 2016; IPBES 2019). Introduced species become invasive when they successfully overcome barriers related to the four different stages of the invasion process (i.e., transport, introduction, establishment, and spread; Colautti et al. 2004; Blackburn et al. 2011). Factors contributing to invasion success, including introduction effort (number of founders and/or introduction events), habitat, and species characteristics, can differentially impact the probability of a species successfully overcoming each stage (Blackburn et al. 2011). The introduction of many individuals or founders can lead to a reduction of extinction probability and can enable populations to more readily adapt to new environments (Blackburn et al. 2015). As well, multiple introduction events can increase the probability of genomic admixture and can result in increased fitness (Keller and Taylor 2010; Basso et al. 2019). Furthermore, habitat characteristics influence invasion success (Lodge 1993; Kolar and Lodge 2001) as they affect the adaptive potential of the introduced species (Le Roux et al. 2008; Vahsen et al. 2018). Additionally, different species characteristics could promote expansion success: for example, explorative and dispersive behaviour may affect the survival of introduced individuals (Holway and Suarez 1999; Chapple et al. 2012; Carere and Gherardi 2013), and a generalist diet could increase survival as more resources are available for generalist species in the invasive range (Fisher and Owens 2004). Life history traits influencing population growth are particularly important at two different stages: 1) the introduction stage because stochastic events can lead to higher risks of extinction, suggesting that smaller founding populations with slow population growth may be more susceptible to extinction (Capellini et al. 2015); and 2) the spread stage because population growth is essential when dispersing individuals form new populations in their introduced range (Capellini et al. 2015).

Life history strategies of species can be placed along a “fast-slow continuum” (Stearns 1983; Promislow and Harvey 1990; Dobson and Oli 2007a, but see Bielby et al. 2007). Fast species –species with life history traits that accompany rapid population growth, such as early maturity and frequent and large litters– are generally short-lived with greater fecundity, whereas species on the slow end of the continuum are long-lived with lower fecundity (Stearns 1983; Promislow and Harvey 1990; Dobson and Oli 2007b). Nonetheless, species within orders or even populations within species can show variation along this continuum (Dobson and Oli 2007a,b). Bat species, for example, may show a combination of life history traits either related to a fast or slow life history (Culina et al. 2019). In invasion ecology, fast life history traits are suggested to promote the population growth and spread of introduced species, thus favouring species ‘invasiveness’: for example, success of mammals at the introduction stage is related to having large, more frequent litters and a longer reproductive lifespan, while establishment success is related to larger litters, longer reproductive lifespan and greater introduction effort, and their spread success is associated with larger, more frequent litters and higher introduction effort (Capellini et al. 2015). A fast life history influences not only invasion success in mammals (Capellini et al. 2015, but see Sol et al. 2008), but also in reptiles, amphibians (van Wilgen and Richardson 2012; Allen et al. 2017), and fishes (Liu et al. 2017). In birds, on the contrary, a slow life history increases the potential to be a successful invader (Sæther et al. 2004; Sol et al. 2012). Although, some very successful invasive birds do not have a slow life history (Markula et al. 2016). Thus, for different species, diverse life history traits may influence their potential for invasion success and exceptions to the general trends occur in all animal taxa.

Nevertheless, introduction effort is the major factor amongst those affecting invasion success (Lockwood et al. 2005; Colautti et al. 2006; Simberloff 2009; Blackburn et al. 2015). Thus, both fast life history traits and a high introduction effort contribute to the probability of established species to successfully spread in the new range (Capellini et al. 2015; Allen et al. 2017). However, the role of introduction effort in explaining species invasiveness could be difficult to interpret in certain taxonomic groups, such as squirrels, as some species are known to be successful invaders without a high introduction effort (either number of founders, introduction events, or both). For example, in 71.4% of successful invasions of grey squirrels, *Sciurus carolinensis*, the number of founders did not explain invasion success because fewer than ten individuals were released (Wood et al. 2007; Lawton et al. 2010). Also in the Pallas’s squirrel, *Callosciurus erythraeus*, and the Finlayson’s squirrel, *Callosciurus finlaysonii*, the number of founders did not prevent them from establishing in their introduced ranges, even though fewer than seven individuals were released (Bertolino 2009; Bertolino and Lurz 2013; Benitez et al. 2013). In the case of the Siberian chipmunk, *Eutamias sibiricus*, some invasive populations were founded by fewer than 10 (Mori et al. 2018b), although some of those have multiple origins (Mori et al. 2018a).

We used Barbary ground squirrels, *Atlantoxerus getulus*, as an exemplar of a species whose introduction effort was extremely low, as only 2 or 3 individuals from one source population in Morocco were introduced to the island of Fuerteventura, Canary Islands, Spain in 1965 (Fig. 1; Machado 1979; Kratzer et al. 2020). In 37 years, the Barbary ground squirrel spread across the entire island, favoured by five translocation accounts from the original founder locality to new localities on the island (Machado 1979; Machado and Domínguez 1982; López-Darias, unpubl. data). Since the introduction, their population has grown to an estimated one million animals (López-Darias 2007), despite low genetic diversity and a small effective population size of 77 individuals (Kratzer et al. 2020). The species has negative ecological, economical, and public health impacts. They consume native and endemic snails (Machado and Domínguez 1982; Groh and García 2004; López-Darias 2007), prey upon some critically endangered species (Bañares et al. 2003), and feed upon the eggs of small native and endemic passerine birds (López-Darias 2007). Their ecological impacts go beyond direct predation upon species, as they alter plant-animal interactions of fleshy-fruited plant species and ruderal weeds, including herbaceous plants of native or introduced origin (López-Darias and Nogales 2008, 2016). Competition with the endemic shrew, *Crocidura canariensis,* listed as vulnerable by the IUCN (Hutterer 2008), is suggested (López-Darias 2008; Traveset et al. 2009), but no quantitative and conclusive data are available. Additionally, these ground squirrels carry parasites that impact native fauna as well as public health (Lorenzo-Morales et al. 2007; López-Darias et al. 2008b), and cause damage to agricultural activities (Machado and Domínguez 1982; López-Darias 2007). Moreover, these ground squirrels alter the population dynamics of predators in two ways (López-Darias 2007). First, the ground squirrels are the main prey of an endemic subspecies of Eurasian buzzard, *Buteo buteo insularum,* whose population has increased on Fuerteventura in the last 40 years, possibly because they prey upon this abundant, new prey (Gangoso et al. 2006; López-Darias 2007). Second, the ground squirrels positively impact invasive feral cat populations (Medina et al. 2008). Despite the notable invasion success of the species on the island, nothing is known about the life history traits that could have influenced their population growth.

**Fig. 1.**
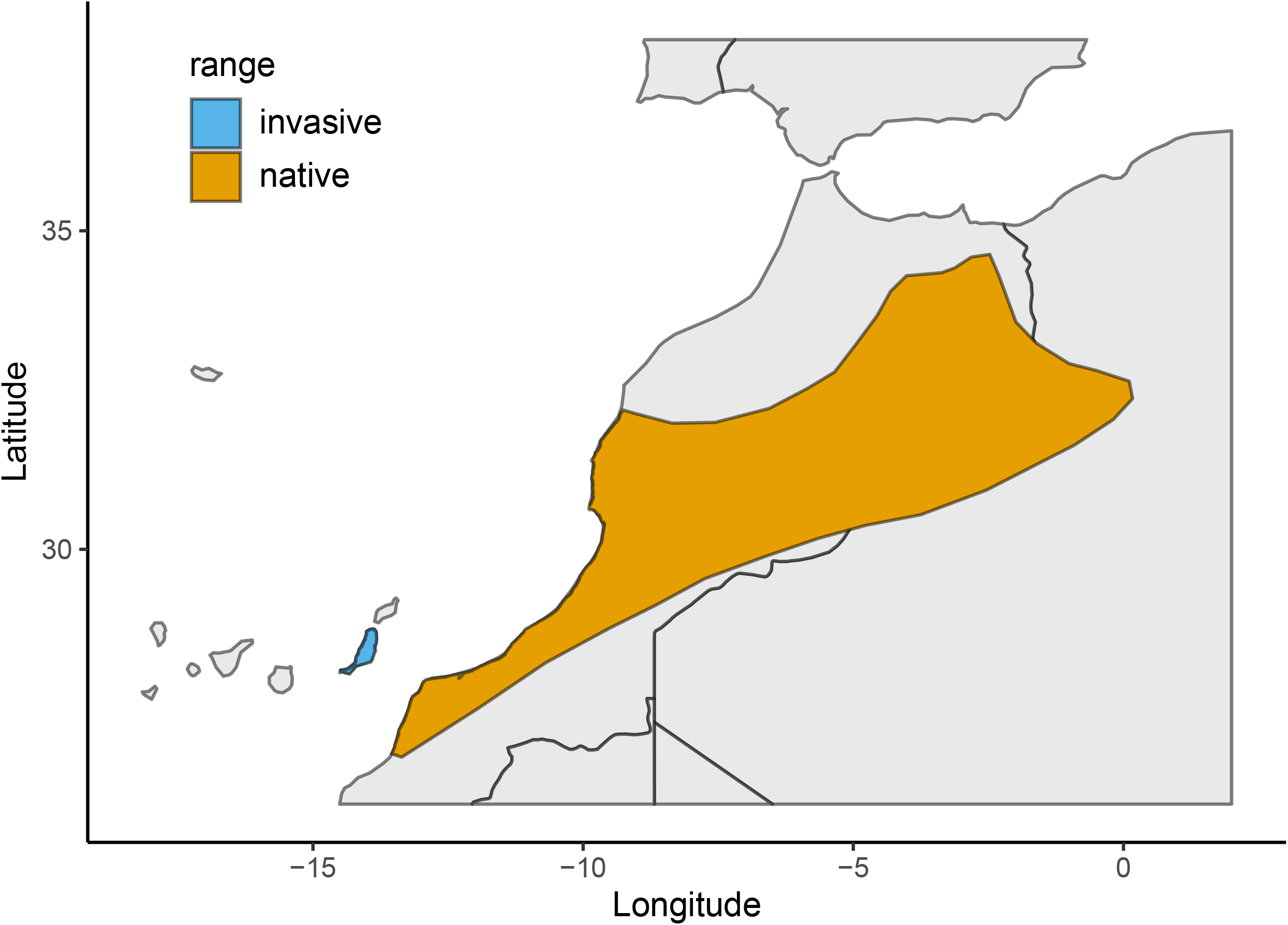
The native distribution in Morocco and the introduced distribution (Fuerteventura, Canary Islands) of Barbary ground squirrels, *Atlantoxerus getulus*. Distribution map derived from the IUCN Red List Mammals (IUCN 2008)

The objective of this manuscript is threefold. First, we investigated the life history traits of the Barbary ground squirrel, a species with a very low introduction effort (Machado 1979; Kratzer et al. 2020). Second, we studied whether the Barbary ground squirrel’s exceptional invasion success was due to a very fast life history profile through a comparison of these traits to other successfully invaded mammals using the dataset provided by Capellini et al. (2015). Third, we examined whether number of founders and/or a fast life history influence invasion success of squirrels using the dataset provided by Bertolino (2009). Our results will be essential to set up effective management strategies for Barbary ground squirrels, as knowing life history traits, such as when females come into estrus or if they can breed year-round, is essential to better plan population control programs. Knowing the life history traits will also improve potential spatially explicit population models, which will be useful for future conservation programs (Bertolino et al. 2020). In a broader comparative context, our results contribute to our understanding of invasion success when introduction efforts are minimal, especially in squirrels.

## Methods

### Life history traits of invasive Barbary ground squirrels

Of the Canary Islands’ seven main volcanic islands, Fuerteventura is the closest to the African continent (approximately 100 km away), the oldest, and has the second-lowest altitude (807 m a.s.l.). The island has an arid climate – high temperatures (≈ 20 °C) and low rainfall (on average < 100 mm/year) – characterised by trade winds, which have resulted in a semi-desert habitat (López-Darias and Lobo 2008). We studied the invasive population of Barbary ground squirrels in the northwest of Fuerteventura (28°34’60” N, 13°58’0” W). Our study site was characterised by stony plains with xerophytic scrubland, interspersed with ravines caused by erosion and abandoned cultivated areas that are fenced by man-made rock walls and dams.

The semifossorial Barbary ground squirrel is a social species: adult females share sleeping burrows with other related females, and males share sleeping burrows with unrelated males and subadults of either sex (van der Marel et al. 2020). We defined these sleeping burrow associations as social groups (van der Marel et al. 2020). Throughout the day, both male and female groups can be active in the same area (van der Marel et al. 2020).

To determine the life history traits of the species, we trapped Barbary ground squirrels using live traps (Tomahawk Co., WI, USA 13 × 13 × 40 cm) baited with peanut butter during three consecutive years (February through July 2014, January through July 2015, and January through June 2016) following the procedures described in van der Marel et al. (2019, 2020). Upon capture, we transferred squirrels to a conical handling bag (Koprowski 2002), where every adult squirrel received a passive integrated transponder (PIT) tag (Avid Identification Systems, Inc., Norco, CA, USA) for individual identification.

We first assessed whether there was a distinct breeding season in the Barbary ground squirrels by regularly trapping and observing male and female reproductive progress. We reported the reproductive status of adult males through the year from non-scrotal to descending scrotum, fully scrotal, and regressing scrotum. Subadult males were those individuals that have reached adult body size, were over six months old, but did not have descended testes (Waterman 1996). Female subadults were individuals that have reached adult body size, were over six months of age, but their vulva and nipples were not swollen during the mating season (indicative of no reproduction). We assessed the reproductive progress of adult females by measuring gestation length from oestrus date until parturition date. We estimated the day of oestrus following previous recommendations by Pettitt et al. (2008) and Waterman (1996). Lactation length was measured from the day of parturition to juvenile emergence. We determined the parturition date by trapping and weighing the females more extensively around their expected parturition date and by palpating their lower abdomen for embryos. The nipples of immature (subadult) females are small but after their first successful parturition, the nipples elongate and swell during lactation. Large bite marks surrounding the nipples indicated the weaning period. We calculated the weaning age from the day of juvenile emergence until the bite marks around the nipples of the mothers started to subside. After weaning, the nipples of adult females stay elongated but not swollen (Waterman 1996). We estimated age at first birth in females with a known date of birth. As our sample size was small, we also estimated age at first birth in females with an unknown day of birth using the known date of emergence from their natal nest burrows. For these females, we added the average lactation length to the date of emergence to estimate their age at first birth. We only used females, because we could be certain they had mated due to the swelling of their vulva and presence of copulatory plugs. As age at first birth violated the assumptions of normality and homoscedasticity of a parametric test, we tested for a difference in our two estimates (females with a known date of birth *vs.* females with a known date of juvenile emergence) using a Mann-Whitney *U* test.

To locate nest burrows of adult pregnant females, we fitted them with radio collars (3.6 g PD-2C transmitters, Holohil Systems Ltd., Carp, Ontario, Canada) just prior to juvenile emergence. We radiotracked the females using an R-1000 telemetry receiver (Communications Specialists, Inc., Orange, CA, USA) and a Yagi antenna model F150-3FB (Advanced Telemetry Systems, Inc., Isanti, MN, USA). Upon juvenile emergence, we extensively trapped at nest burrows to sex, measure, and mark the juveniles (individuals newly emerged from their nest burrow up to six months of age) with ear tags (#1005 Size 1 Monel, National Band and Tag Co., Newport, KY, USA). We also determined litters per year and litter size. We used a Kruskal-Wallis test to analyse differences in average litter size across years. We calculated the litter sex ratio as the proportion of male juveniles out of the total emerged juveniles (Charnov 1982; Ryan et al. 2012), which we analysed using a chi-square test of independence. We tested whether litter sex ratio differed from parity using an exact binomial test. Using a chi-square test of independence, we analysed differences in juvenile survival rate, measured as the proportion of emerged juveniles that survived to one month. Occasionally, mothers merged their litters with the litters of other adult females before all juveniles were trapped and marked, resulting in unmarked juveniles at a communal burrow. These litters were excluded because we were uncertain of the identity of the mothers. Finally, we calculated maternal success rate as the proportion of females successfully raising their litter to emergence and used a chi-square test of independence to test for differences in maternal success rate.

We were unable to measure neonatal body mass in the field because the females give birth within their nest burrow; therefore we used neonatal body mass recorded during an earlier study with the Barbary ground squirrels on Fuerteventura (mean ± SE = 8.1 ± 0.5 g, *n* = 5 juveniles; Machado and Domínguez 1982). We measured adult body mass using a spring scale (± 5 g; Pesola AG, Baar, Switzerland) and focused on masses outside of the reproductive period, selecting males that were non-scrotal and females prior to the breeding season or after their offspring were weaned. We measured average adult lifespan for individuals of known age and for individuals of uncertain age (individuals that did not survive into adulthood were excluded). We tested for sex differences in average lifespan using Mann-Whitney *U* tests. We calculated adult survival as the proportion of adults in a group that were trapped in the following field season (January 2015, 2016, and 2017). We censored adult survival for squirrels that were already adults in our first field season and were still alive in our last field season, i.e., squirrels with uncertain age. We performed the Kaplan-Meier approach on our censored data using the ‘Survival’ package version 2.43-1 (Therneau and Grambsch 2000; Therneau 2015). Reproductive lifespan was calculated from the maximum lifespan/longevity (5.0 year) reported in Aulagnier et al. (2013), transformed into days and age at first birth in days following Capellini et al. (2015).

To understand the contribution of life history traits and introduction effort to invasion success, we questioned whether the Barbary ground squirrel’s exceptional invasion success (despite low introduction effort) is due to a very fast life history profile. We compared the life history profile of the Barbary ground squirrel population to the life histories of other invasive mammals and other invasive squirrels using the dataset provided by Capellini et al. (2015). These authors derived the life history data from the panTHERIA database (Jones et al. 2009), where the maximum lifespan was provided for individuals in captivity, while data in the wild exist for three out of the five introduced squirrel species. Therefore, we changed the maximum lifespan of gray squirrels to 12.5, red squirrels to 7 (Barkalow and Soots 1975) and Siberian chipmunks to 6 years (Chapuis et al. 2011). As reproductive lifespan is derived from longevity, we also changed those values for the species in question. We performed all the statistical analyses with alpha set to 0.05 in R version 3.5.1 (R Core Team 2017). We reported the mean ± standard error (*SE*).

### Life history vs. introduction effort in explaining the invasion success of squirrels

We used a Bayesian Phylogenetic Multilevel Model to test the impact of life history traits (longevity, body mass, gestation length, weaning age, litter size, litters per year, age at first birth, and reproductive lifespan) and introduction effort (number of founders and times introduced) on the invasion status of squirrels (Bertolino 2009). For our phylogeny, we selected the squirrel species reported in Bertolino (2009) from a mammalian supertree provided by Rolland et al. (2014). In our model, we used invasion status (successfully spread or not; Bernoulli distribution) as our dependent variable. We tested for collinearity between our variables using the variance inflation factor function from the ‘CAR’ package version 3.0-2 (Fox and Weisberg 2011) and excluded covariates with a variance inflation factor value above three. We included the number of founders, weaning age, litters per year, and age at first birth as our covariates (ESM 1) and standardised these variables. We used species as a random factor to account for any effects that are independent of the phylogenetic relationship between species as we had multiple introduction events per species (n = 8 species ranging between 1 and 28 introduction events with 878 data points in total, Bertolino 2009). We also accounted for the phylogenetic signal between species. We ran two models, one without added variability of the independent variables within species and one with it. As only the number of founders was variable per species, we only had to account for this covariate. We used the ‘get_prior’ function to set our prior. As settings for our model, we used three chains, two cores, 10000 iterations, and 1500 as the warmup (burnin), and we set the ‘adapt_delta’ to 0.99. We visually inspected for model convergence and we assessed mixing by ensuring that the effective sample sizes for all parameters were above 1000. We selected the model with the lowest loo information criterion (Vehtari et al. 2017). We assessed the influence of each independent variable on invasion status by analyzing whether the posterior distribution crosses zero. If the posterior distribution of the parameter estimates (β) crosses zero then that variable has a negligible effect on the dependent variable, whereas if it substantially shifted from zero then that variable has an influential effect. For this analysis, we used the R package ‘brms’ (Bürkner 2017, 2018) from the Stan environment (http://mc-stan.org/). We performed the statistical analyses in R version 4.0.1 (R Core Team 2020) and provided the R code on a Github repository (https://github.com/annemarievdmarel/lifehistory_invasion; van der Marel 2020).

## Results

### Life history traits of invasive Barbary ground squirrels

Reproduction in Barbary ground squirrels was seasonal; in 2014, mating started at the end of January and ended mid-March (reproductive progress observed for 32 males and 40 females), whereas in 2015 (*n* = 52 males and 69 females) and 2016 (*n* = 55 males and 82 females) mating started at the beginning of January and late December and ended mid-February and at the end of January, respectively. Males were scrotal for 76.3 ± 4.0 days (*n* = 65) after the last day of the mating period, except for 8 subadult males in 2014 and one in 2016. It took adult males on average 152.1 ± 5.0 days (*n* = 54) to fully regress their scrota. Within each social group, all adult females bred. Gestation length was 43.5 ± 0.4 days (range 39-49 days; *n* = 47 females) and lactation lasted 42.3 ± 0.5 days (range 39-46 days; *n* = 43 females). Most females produced their first litter when they are approximately one year old, except for the six subadults in 2014. The estimates of age at first birth (primiparity) calculated for females with a known date of birth (343.4 ± 11.0 days, *n* = 8) and for females with a known date of emergence (345.5 ± 5.7 days, *n* = 15) did not differ (Mann-Whitney *U* test: *U* = 53, *P* = 0.67). Therefore, the average age of first birth was 344.8 ± 5.2 days (*n* = 23 females).

Over two years (2015 and 2016), 15 of 138 pregnancies failed (10.9%). Females rebred in 11 of these failed pregnancies (73.3%), although five of these later pregnancies failed again (45.5%). Maternal success rate (proportion of females that successfully raised their litter to emergence) did not differ among years (2014: 0.78, 2015: 0.76, and 2016: 0.79; chi-square test of independence: χ^2^_2_ = 0.71, *P* = 0.70). In 2016, eight females of a total of 80 (10.0%) were observed to have a second litter in one breeding season after they had successfully bred once, but we do not have data on their litters because we left the field before the emergence of these late litters. In 2014, 112 juveniles emerged from 28 litters; in 2015, 161 juveniles from 44 litters; and in 2016, 166 juveniles from 63 litters. Average litter size did not differ among years (2014: 2.3 ± 0.3, 2015: 3.4 ± 0.4, and 2016: 3.1 ± 0.2; Kruskal-Wallis test: *H*_2_ = 3.91, *P* = 0.14). The mean percentage of adult females that were successful in raising one litter during the three years was 75.5 ± 2.0%, with a litter size of 3.0 ± 0.2 [1 – 8] juveniles per litter (*n* = 62 litters). Litter sex ratio was 0.50 male across all years and did not differ among years (2014: 0.50, 2015: 0.47, and 2016: 0.52; chi-square test of independence: χ^2^_2_ = 0.24, *P* = 0.89), nor did it differ from parity (exact binomial test: two-tailed *P* = 1.0, 95%-CI = [0.42 − 0.59], *n* = 45 litters from 40 distinct females). Juvenile mortality was significantly lower in 2014 (19.4%) compared to 2015 (50.3%) and 2016 (49.4%), respectively (χ^2^_2_ = 31.24, *P* < 0.001).

Average adult body mass outside of the reproductive season was 221.1 ± 5.7 g (*n* = 116 individuals) with males weighing 227.0 ± 6.4 g (*n* = 51), and females weighing 204.0 ± 5.0 g (*n* = 65). As average lifespan did not differ between adult males and females (lifespan of individuals of known age, Mann-Whitney *U* test: *U* = 247.5, *P* = 0.20; and lifespan of individuals of uncertain age included, *U* = 10187, *P* = 0.95), we combined the sexes. The estimated average lifespan of the Barbary ground squirrel in our sites calculated over three years was 1.48 ± 0.09 years of age for individuals with known age (*n* = 52), but 1.76 ± 0.06 years when individuals with uncertain age were included (*n* = 239). The survival rate did not differ between males and females (*P* = 0.41) and was 0.89 ± 0.02, 0.79 ± 0.03, and 0.73 ± 0.04 at age 1, 2, and 3 years, respectively. Reproductive lifespan calculated using the maximum lifespan/longevity of five years (Aulagnier et al., 2013) was 1480.2 days (4.1 years).

The life history traits of Barbary ground squirrels were all lower compared to the traits of other invasive mammals, except for litter size, weaning age and litters per year (Table 1, Fig. 2).

**Table 1.** Life history traits median values and sample size (*N*) for all mammals, the species of the family Sciuridae (squirrels, including Barbary ground squirrels *A. getulus*) that successfully spread in their introduced range (invasive species), and the Barbary ground squirrels only

|  | Mammals<sup>a</sup> |  | Only squirrels<sup>a</sup> |  | *A. getulus* only |  |
| --- | --- | --- | --- | --- | --- | --- |
|  | *N* | median | *N* | median | *N* | median |
| Introduction effort | 47 | 26 | 6 | 11 | 1 | 1 |
| Body mass (g) | 47 | 6361.6 | 5 | 349.6 | 1 | 220 |
| Neonatal body mass (g) | 47 | 113.0 | 4 | 8.8 | 1 | 8.0 |
| Litter size | 47 | 2.3 | 5 | 3.8 | 1 | 3.0 |
| Litters per year | 47 | 1 | 6 | 2 | 1 | 1 |
| Age at first birth (days) | 45 | 403.3 | 5 | 357.5 | 1 | 350.3 |
| Gestation time (days) | 47 | 66.4 | 5 | 41.4 | 1 | 42.0 |
| Weaning age (days) | 45 | 60.9 | 5 | 54.8 | 1 | 67 |
| Longevity (years) | 47 | 20.0 | 5 | 7<sup>b</sup> | 1 | 5 |
| Reproductive lifespan (years) | 45 | 18.3 | 5 | 6.3<sup>b</sup> | 1 | 4.1 |

<sup>a</sup> Life history traits adopted from Capellini et al. (2015), who derived the life history data from the PanTHERIA database (Jones et al. 2009), except for the *A. getulus* data.
<sup>b</sup> In the dataset of Capellini et al. (2015) longevity for the species in captivity was provided if longevity was not known for the species in the wild, which overestimates longevity and the derived reproductive lifespan. We changed the longevity of gray squirrels to 12.5, red squirrels to 7 (Barkalow and Soots 1975) and Siberian chipmunks to 6 years (Chapuis et al. 2011).

**Fig. 2.**
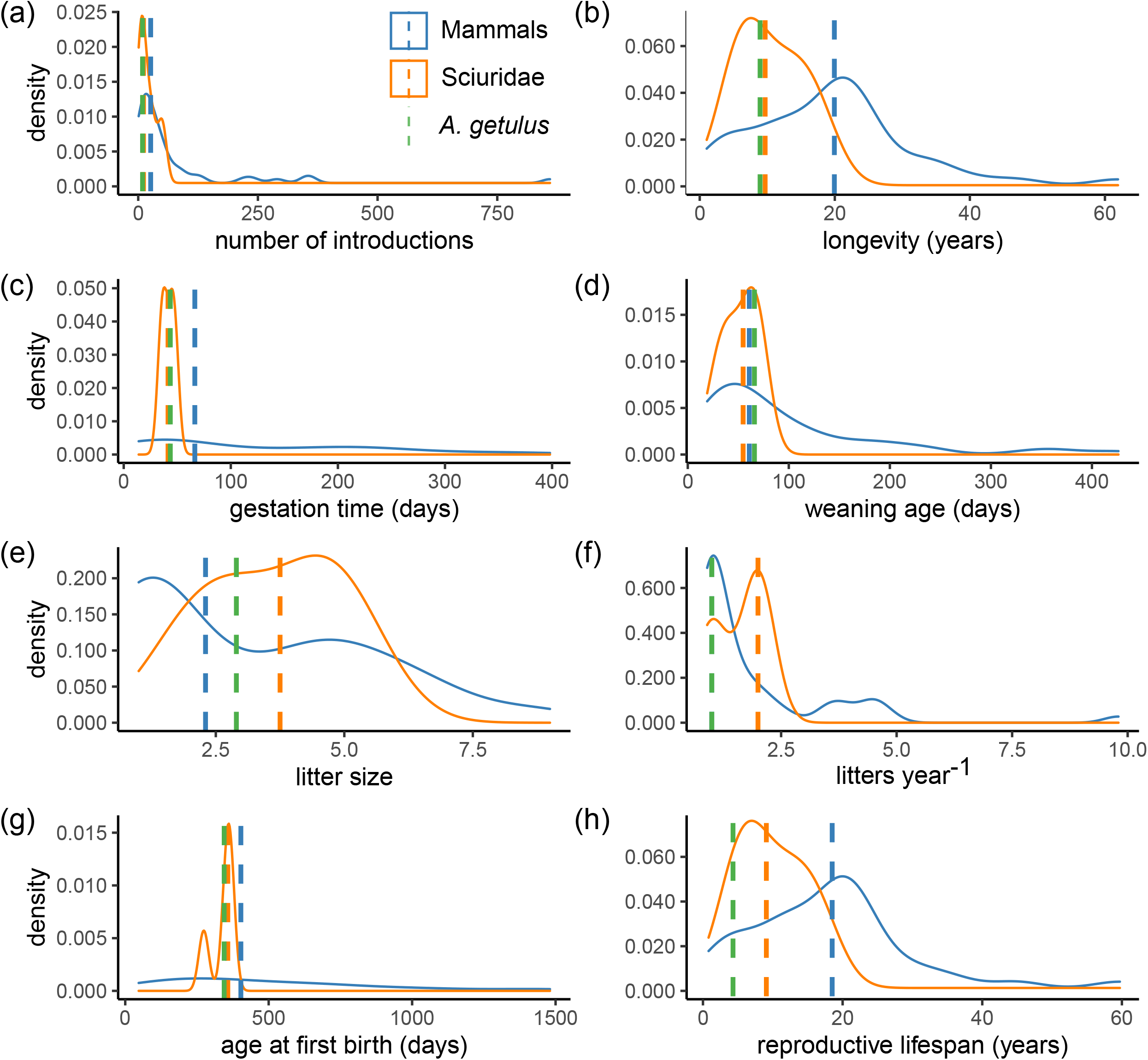
Density plots of a) number of introductions and several life history traits such as b) longevity, c) gestation time, d) weaning age, e) litter size, f) litters per year, g) age at first birth, and h) reproductive lifespan, for all mammals (blue) and all species of the family Sciuridae (including our data for Barbary ground squirrels *Atlantoxerus getulus*) that successfully spread in their introduced range (invasive species) (orange). Dashed lines are the median values for mammals (blue) and squirrels (orange), and the point values for the traits of Barbary ground squirrels (dashed green line)

### Life history vs. introduction effort in explaining the invasion success of squirrels

The model without added variability of the independent variables within species (looic = 71.7) best explained whether squirrels were successful invaders in comparison to the model with added variability of the independent variables within species (looic = 82.6). However, neither number of founders, nor life history characteristics explained invasion status of introduced squirrels (Fig. 3), as all the posterior distributions of the covariates crossed zero (Fig. 4). We found effects that were both dependent and independent of the phylogenetic relationship between species (random effect species: 95% CI = [0.01 – 1.99] and phylogenetic signal: 95% CI = [0.04 – 4.13]).

**Fig. 3.**
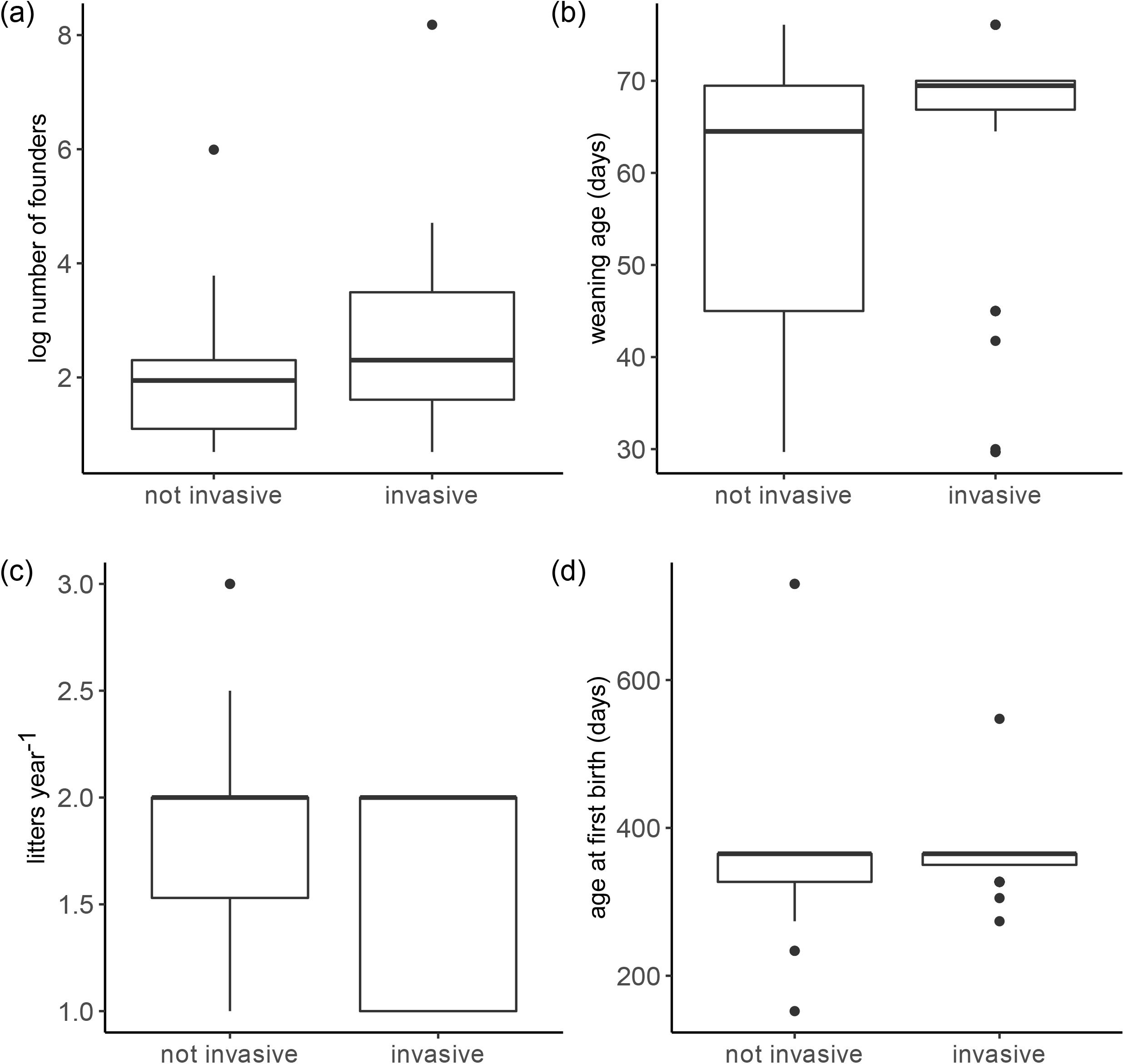
The (a) log number of founders, (b) weaning age, (c) litters per year, and (d) age at first birth for squirrel species that did not successfully spread (not invasive) and that did successfully spread (invasive) into their new geographical range. The dark line is the median, the box edges are the upper and lower quartiles, the whiskers are 50% from the median, and the closed circles are the outliers, calculated as the values smaller or larger than 1.5 times the box length (i.e., upper-lower quantile). Note the non-standardized y-values

**Fig. 4.**
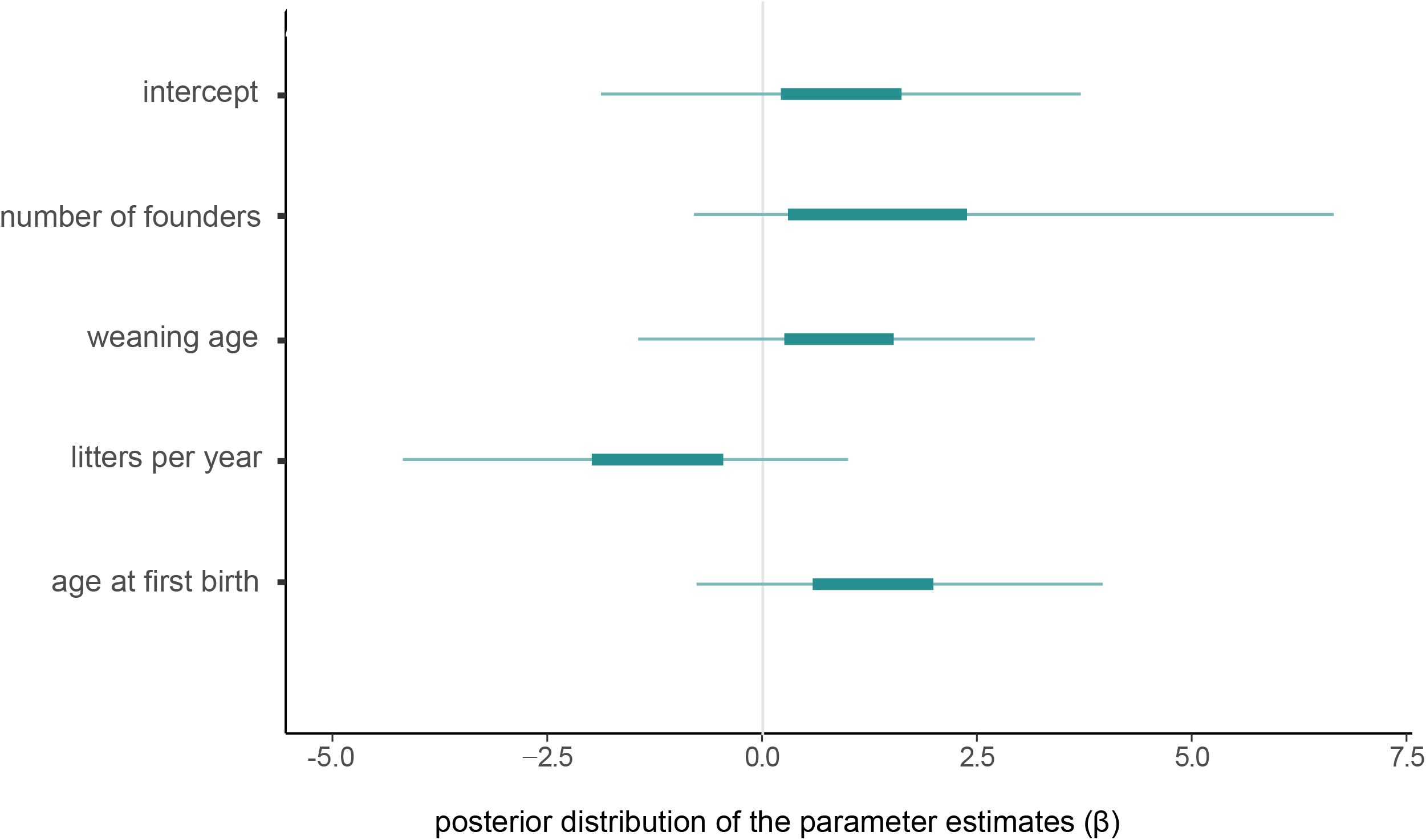
The posterior uncertainty intervals of the parameter estimates (β) of introduction effort and life history traits. The thick segments are the 50% intervals, and the thinner outer lines are the 95% intervals. The distributions that cross zero have a negligible effect on invasion status, whereas distributions that do not contain zero have a strong effect on invasion status. Plot designed using the ‘bayesplot’ package (Gabry et al. 2019)

## Discussion

Our study provides the first detailed data on the life history and population dynamics of an invasive population of Barbary ground squirrels, a population with one of the lowest introduction efforts (one pair or three individuals as founders from one source location, Kratzer et al. 2020), but a great invasion success (López-Darias 2007). The Barbary ground squirrel represents the fast end of the “fast-slow continuum” of life history traits, because they are small-bodied (Machado and Domínguez 1982, this study), mature within their first year and females have large and frequent litters. Barbary ground squirrels’ exceptional invasion success (despite low introduction effort, Kratzer et al. 2020) is not due to a remarkable fast life history profile, as their life history is comparable to other invasive species that do not have such a low introduction effort. When we analysed the effects of introduction effort and life history traits further, we found that neither number of founders nor life history traits influenced invasion status of introduced squirrel species. These results suggest that especially for squirrels, other traits besides introduction effort and life history traits influence their probability to successfully invade a new range (e.g., Signorile et al. 2014; Mori et al. 2018b), which we discuss further below.

### Potential factors influencing invasion success of Barbary ground squirrels

The Barbary ground squirrel can be considered a genetic paradox of invasion (Estoup et al. 2016). The invaded population was founded by 2 or 3 individuals, which has resulted in a low genetic diversity and a high level of inbreeding in the invasive population (Kratzer et al. 2020). A high degree of inbreeding could lead to inbreeding depression, i.e., a high impact of deleterious alleles on the average fitness of offspring (Ralls et al. 1979; Ralls and Ballou 1982). However, in this particular case of the Barbary ground squirrel, with a population in the introduced range estimated at around one million individuals (López-Darias 2007), argues against inbreeding depression, because this population has successfully reproduced and grown. Although we have shown that the squirrels have a fast life history, this factor alone could not explain their invasion success. Besides a fast life history, other aspects, such as anthropogenic, habitat or species-level factors, may have favoured their invasion success. First, multiple anthropogenic translocations of the initial founder population (Machado 1979; Machado and Domínguez 1982; López-Darias, unpubl. data) may have had a significant effect on the invasion success of the Barbary ground squirrel.

Second, climatically matched habitat characteristics could impact the adaptive potential and population growth of an introduced species in their new range (Lodge 1993; Kolar and Lodge 2001; Forsyth et al. 2004; Hayes and Barry 2008). For Barbary ground squirrels, climate conditions of the source location (Sidi Ifni, Morocco, Machado 1979; 133 mm rainfall/year and 19.2 °C, Merkel, 2019) are very similar to the habitat on Fuerteventura (Machado 1979; <100 mm rainfall/year and ~ 20 °C, López-Darias and Lobo 2008). Since the climate is so similar, it is no surprise that climate did not affect the squirrel’s abundance in their invasive range (López-Darias and Lobo 2008). As well, habitat preferences are similar between the native (Rihane et al. 2019) and the invasive range (López-Darias and Lobo 2008). Moreover, Barbary ground squirrels are reported to have a low diversity of parasites (López-Darias et al. 2008b), and a lower number of predator species (species richness) in the invasive range (Gangoso et al. 2006; Medina et al. 2008) compared to the native range (Machado 1979). Additionally, the main current aerial predator (the Eurasian buzzard) had an atypically small population at the time the squirrels were introduced (Gangoso et al. 2006), suggesting a release from predation around the establishment stage of Barbary ground squirrels. Only 2% of the diet of the only terrestrial predator of the Barbary ground squirrel, feral cats, consists of the squirrels (Medina et al. 2008). Nevertheless, predation pressure may still be significant if the number of feral cats is large. No information regarding the population size of feral cats is available, but cats were introduced to the Canarian archipelago in the 15^th^ century and are now present in every habitat of each main island (Medina et al. 2008). Feral cats are not only potential predators of the squirrels but they are also a problem for wildlife conservation, especially on islands (Medina et al. 2011; Doherty et al. 2016). Overall, Fuerteventura can be considered a suitable environment for the Barbary ground squirrel (López-Darias et al. 2008a).

Barbary ground squirrels also may have become successful invaders due to behavioural advantages, as favourable behavioural traits promote success at each stage of the invasion (Holway and Suarez 1999; Chapple et al. 2012; Carere and Gherardi 2013). Species with increased interspecific or decreased intraspecific aggression, and behaviours enhancing their dispersal, e.g., exploration, activity, and boldness, should perform better in their new habitat (Holway and Suarez 1999). These behavioural traits have been hypothesised to represent an invasion syndrome (Sih et al. 2004; Chapple et al. 2012), and can be linked to life history traits, which then result in “pace-of-life” syndromes (Réale et al. 2010). Often species with a fast pace-of-life syndrome, i.e., more explorative and bold species, have lower survival chances because they are more exposed to predators and parasites (Réale et al. 2010). For introduced species, a fast pace-of-life may be advantageous since fewer predators and parasites result in higher survival chances. In Barbary ground squirrels, this hypothesis is supported, because more explorative individuals are better at avoiding predation, resulting in greater survival chances (Piquet et al. 2018).

To conclude, generalist species – species not specialised in habitat use or diet – are suggested to better adapt to a variable environment and to have a better chance of becoming successful than specialist species (Fisher and Owens 2004). The Barbary ground squirrel has a generalist diet because they are omnivorous and eat not only seeds, nuts, and fruits, but also native mollusks (Machado and Domínguez 1982; López-Darias and Nogales 2008). Additionally, we have recorded them feeding on the horns and hooves of goat carcasses (van der Marel, pers. obs.), supporting the suggestion that the squirrels could be mineral-limited (Machado 1979). A generalist diet could have helped the squirrels survive dry years with scarce resources, as they would have a wider niche breadth and not depend on a limiting resource. For example, in Darwin’s tree finches, generalist species have a more varied diet in dry years compared to wet years (Christensen and Kleindorfer 2009). Thus, multiple different aspects could have helped the squirrels overcome barriers in their introduced range.

### Life history vs. introduction effort in explaining the invasion success of squirrels

Neither introduction effort measured as the number of founders nor life history traits affected the invasion status of squirrels as all the posterior distributions of the covariates crossed zero. The life history traits may not be varied enough among the related squirrel species studied here to show any effects on invasion status, as we do see an effect of phylogeny in our model. It seems counterintuitive that introduction effort does not impact invasion success, because multiple advantages are associated with a high introduction effort (Lockwood et al. 2005; Colautti et al. 2006; Simberloff 2009; Blackburn et al. 2015).

First, multiple introduction events increase fitness (Keller and Taylor 2010) and eventually genetic diversity (Dlugosch and Parker 2008). Genetic diversity loss is U-shaped when plotted against time since first introduction, suggesting that an invasive population regains its genetic diversity over time, which can be promoted when multiple introduced populations connect (Dlugosch and Parker 2008).

Second, more founders reduce the risk of extinction and promote adaptability to the new environment through increased genetic diversity (Dlugosch and Parker 2008; Blackburn et al. 2015). For example, lower genetic diversity caused by a small number of founders reduced the spread rate of grey squirrels (Signorile et al. 2014). Still, the species did successfully spread despite a low introduction effort, albeit slower, suggesting that a loss in genetic diversity does not imply a loss in the adaptive potential of an introduced species *per se* (Dlugosch and Parker 2008). Instead, the probability for population growth from a small founding population in tree squirrels may be attributed to a high reproductive output (Wood et al. 2007).

Additionally, climate and habitat suitability may be a better predictor for invasion success than the number of founders or a fast life history in squirrels, similar to mammals introduced to Australia (Forsyth et al. 2004). Normally, demographic stochasticity negatively affects population growth in small populations; however, other factors unrelated to demographic stochasticity, such as unsuitable habitat or climate, may negatively impact large populations as well (Blackburn et al. 2015). As such, small populations may then persist when other factors, such as climate and habitat suitability, work instead in their favour.

Also, other factors may help species become invasive with low introduction effort. For example, the invasion success of Siberian chipmunks is influenced by food provisioning by humans, especially in urban environments, and the absence of competitors at the time of introduction (Mori et al. 2019). Thus, for squirrel species particularly, other factors – climate and habitat suitability, enemy release, a fast life history – can be better predictors of invasion success than introduction effort.

### Concluding remarks and future studies

Barbary ground squirrels have a fast life history strategy resulting in rapid population growth. If these life history traits have not changed since 1965, their fast life history may have contributed to their invasion success with an extremely low introduction effort on Fuerteventura Island, together with favourable behavioural traits, a generalist diet and good resources, enemy release, and similar habitats and climate. Only by understanding their basic biology will we be in the position to control and minimise the ecological damage this species causes in their new habitat. For a future study, we could run a population model to predict how long it would take for a pair of squirrels to spread across the island or to evaluate the necessary effort to control the population or reduce the number of squirrels on Fuerteventura, while taking the factors influencing their invasion success into account (Merrick and Koprowski 2017; Bertolino et al. 2020).

Our study also aids in constructing a comprehensive framework on the factors, including life history traits and introduction effort, influencing invasion success in mammals. For introduced squirrels, suitable climate or habitat may be more important predictors whether a species becomes a successful invader than introduction effort or life history traits alone. Our results contradict the concept that introduction effort is the key factor influencing invasion success (Lockwood et al. 2005; Simberloff 2009); instead, we argue that the invasion success can be influenced by multiple aspects of the new habitat or the biology of the introduced species.

## Acknowledgements

We thank the owners of our study sites for access to their lands, the Cabildo of Fuerteventura for access at the Estación Biológica de la Oliva, and the IPNA-CSIC for logistical support. We thank E. Anjos for providing useful feedback.

## Declarations

### Ethics approval

All procedures conformed to the guidelines of the American Society of Mammalogists for the use of wild mammals in research, were approved by the University of Manitoba Animal Care and Use Committee (protocol #F14-032) and were permitted by the government of Fuerteventura (Cabildo Insular de Fuerteventura #14885).

### Consent to participate

Not applicable

### Consent for publication

All authors whose names appear on the submission approved the version to be published.

### Availability of data and material

The datasets analysed during the current study are available in the Github repository, https://github.com/annemarievdmarel/lifehistory_invasion (DOI: 10.5281/zenodo.4136880).

### Code availability

The code used during the current study are available in the Github repository, https://github.com/annemarievdmarel/lifehistory_invasion (DOI: 10.5281/zenodo.4136880).

### Authors’ contributions

AM, JMW and MLD conceived and designed the study and collected the data. AM analyzed the data and wrote the manuscript; JMW and MLD provided editorial advice.

